# The impact of phenotypic heterogeneity of tumour cells on treatment and relapse dynamics

**DOI:** 10.1101/2020.11.27.400838

**Authors:** Michael Raatz, Saumil Shah, Guranda Chitadze, Monika Brüggemann, Arne Traulsen

## Abstract

Intratumour heterogeneity is increasingly recognized as a frequent problem for cancer treatment as it allows for the evolution of resistance against treatment. While cancer genotyping becomes more and more established and allows to determine the genetic heterogeneity, less is known about the phenotypic heterogeneity among cancer cells. We investigate how phenotypic differences can impact the efficiency of therapy options that select on this diversity, compared to therapy options that are independent of the phenotype. We employ the ecological concept of trait distributions and characterize the cancer cell population as a collection of subpopulations that differ in their growth rate. We show in a deterministic model that growth rate-dependent treatment types alter the trait distribution of the cell population, resulting in a delayed relapse compared to a growth rate-independent treatment. Whether the cancer cell population goes extinct or relapse occurs is determined by stochastic dynamics, which we investigate using a stochastic model. Again, we find that relapse is delayed for the growth rate-dependent treatment type, albeit an increased relapse probability, suggesting that slowly growing subpopulations are shielded from extinction. Sequential application of growth rate-dependent and growth rate-independent treatment types can largely increase treatment efficiency and delay relapse. Interestingly, even longer intervals between decisions to change the treatment type may achieve close-to-optimal efficiencies and relapse times. Monitoring patients at regular check-ups may thus provide the temporally resolved guidance to tailor treatments to the changing cancer cell trait distribution and allow clinicians to cope with this dynamic heterogeneity.

**Author summary:** The individual cells within a cancer cell population are not all equal. The heterogeneity among them can strongly affect disease progression and treatment success. Recent diagnostic advances allow measuring how the characteristics of this heterogeneity change over time. To match these advances, we developed deterministic and stochastic trait-based models that capture important characteristics of the intratumour heterogeneity and allow to evaluate different treatment types that either do or do not interact with this heterogeneity. We focus on growth rate as the decisive characteristic of the intratumour heterogeneity. We find that by shifting the trait distribution of the cancer cell population, the growth rate-dependent treatment delays an eventual relapse compared to the growth rate-independent treatment. As a downside, however, we observe a refuge effect where slower-growing subpopulations are less affected by the growth rate-dependent treatment, which may decrease the likelihood of successful therapy. We find that navigating along this trade-off may be achieved by sequentially combining both treatment types, which agrees qualitatively with current clinical practice. Interestingly, even rather large intervals between treatment changes allow for close-to-optimal treatment results, which again hints towards a practical applicability.

## Introduction

Cancers are composed of genetically and phenotypically diverse cell populations [1–6], reflecting evolutionary and ecological processes that occur during cancer progression. So far, this heterogeneity has mostly been attributed to the genomic level, where mutations and chromosomal changes are by now routinely detected by diverse molecular techniques. In addition, there are also non-genetic drivers of heterogeneity, such as epigenetic changes, cell differentiation, stochastic gene expression or effects of the microenvironment [4]. The resulting intratumour heterogeneity has to be taken into account, as it can lead to the treatment of only some subclones [7] or the selection of resistant phenotypes [8], which may explain the frequent relapses in many cancer types. Additionally, incomplete sampling of a heterogeneous tumour may hinder the prediction of disease dynamics [5,9,10]. However, if intratumour heterogeneity is exhaustively determined, it may be used as a prognostic factor and guide treatment decisions [5,11,12]. Currently, a strong focus lies on the genetic components of intratumour heterogeneity, and only recently the phenotypic heterogeneity regained clinical interest, mainly after the rise of targeted therapies, a suitable treatment approach only for defined phenotypes [3]. However, genetic heterogeneity does not necessarily map one-to-one to phenotypic heterogeneity [2], and phenotypic heterogeneity may translate into an incomplete and inhomogeneous response to phenotype-dependent therapies [5]. It is therefore imperative to characterize the phenotypic heterogeneity within tumours as well [13, 14]. Understanding and utilizing the phenotypic heterogeneity of a tumour can be aided by ecology and the concept of traits and trait distributions [15, 16]. In this sense, cancer, an evolutionary disease, can indeed be better understood using ecological concepts [17, 18]. We follow this approach and develop an ecological trait-based model to understand the effects of phenotypic trait heterogeneity on treatment outcomes.

The presence of phenotypic intratumour heterogeneity may require novel treatment strategies. Determining the existence of phenotypic subpopulations that likely will provide resistance against phenotype selective therapies is thus a first useful step to decide against exclusive treatment options that would select for these potentially dangerous subpopulations [5]. Secondly, if such resistant cell types are present in low fractions, there may be mechanisms keeping them low, such as trade-offs between treatment resistance and other traits [19]. These can be used to reduce the amount of unfavourable cancer cell types again after intermittently selecting for them by treatment [14, 17, 20–24]. If there are no mechanisms that can suppress resistance once it evolved, a containment strategy may be applied to prolong the time until treatment failure [25]. Another option is to combine different treatment types to create an evolutionary double bind, wherein one treatment renders the other more effective [17,26]. All these adapted treatment schemes require a regular assessment of tumour cell numbers. Given the recent advances in next-generation sequencing, singlecell RNA sequencing, flow cytometry and imaging, however, also obtaining time series for both genetic and phenotypic heterogeneity may become possible in the near future. Here, we assume that such temporally resolved trait information is available and ask how it could best be exploited to improve treatment outcome.

Particularly in light of different treatment options that exert different selection pressures on different traits, such as chemotherapy or immunotherapy, considering the temporal change of trait distributions may be decisive for treatment evaluation. Even though differences in other functional traits are possible and likely, differences in the growth rates of individual cancer cells may be the most obvious aspect of intratumour heterogeneity and also the most decisive for cancer progression. Diverse growth rate trait distributions have been known for a long time [27], and also recently received theoretical interest [28]. To study how treatment interacts with the growth rate trait distribution, we will investigate how two different treatment types, one that depends on the focal trait and one that is independent of it, result in different treatment outcomes. This allows us to predict how their combination may direct the temporal change of the trait distribution to optimize the treatment effect.

Our approach is motivated by the current treatment of acute lymphoblastic leukaemia, for which chemotherapy is the first-line treatment and usually quickly reduces the density of malignant lymphoblasts below the detection threshold. Frequently, however, a fraction of these malignant cells is not eradicated by the treatment but remains in the body as minimal residual disease that causes relapse during or after therapy [29]. We hypothesize that chemotherapy targets fast dividing cells preferentially, as most chemotherapeutic drugs target cell division and thus lead to higher drug-induced death rates in fast-dividing malignant cells. Thus, chemotherapy may exert a selection pressure on cancer cell’s growth rate, eventually favouring slower cells. Due to their slower growth, these cells will only be present in low abundance in the cancer cell population at the initiation of treatment. Still, they may dominate the population in later stages of treatment due to their lower sensitivity to treatment. If the cancer population is driven to low numbers, it becomes vulnerable to stochastic extinction. These stochastic extinction events will be primarily driven by the traits of the slow-growing subset of cancer cells, which might create a reservoir of cells that are less vulnerable to treatment, and eventually grow again once treatment is terminated and cause a relapse. Accordingly, a current approach for post-chemotherapy relapses is to conduct immunotherapy using the bi-specific monoclonal antibody Blinatumomab [30]. Interestingly, this second treatment type’s action is likely independent of growth rate and therefore presents a treatment that is independent of our focal trait. How these two different treatment types operate and interact is to date empirically unknown and justifies theoretical investigation.

Using growth rates as a focal trait, we will investigate how trait (in-)dependent treatment affects the trait distribution of cancer cells, and further how this trait distribution determines treatment trajectories, relapse dynamics and optimal treatment schemes.

## Methods

### Deterministic model

We model the cancer cell population as a collection of Ω subpopulations of size *x_i_* that differ in their growth rates *r_i_*, (Fig. 1). We assume exponential growth for every subpopulation with growth rates *r_i_* (*i* = 1,..., Ω). Further, we assume that the growth rates of individual subpopulations increase linearly from *r*_min_ = *r*_1_ to *r*_max_ = *r_Ω_* (Tab. 1). Genetic and non-genetic drivers may generate heterogeneity within the cancer cell population that manifests as a broadened trait distribution of growth rates [4, 27]. This allows cells to switch to adjacent subpopulations with different growth rates and maintains the width of the trait distribution. We assume that a cell’s switching rate is proportional to its growth rate. Switching to the next slower subpopulation thus occurs at rate *p_S_ r_i_* and switching to the next faster subpopulation happens at rate *p_F_ r_i_*. We include two different treatment types, one that is growth rate-dependent and one that is growth rate-independent. The growth rate-dependent treatment is motivated by the idea that under chemotherapy, the uptake and action of the therapeutic agent is proportional to the growth rate of the cancer cell. Therefore, the rate at which the chemotherapeutic toxins enter the cell, stop cell proliferation and induce cell death is assumed to be proportional to the cell’s growth rate. The growth rate-dependent treatment thus induces a cancer cell mortality *δr_i_* where *δ* captures the trait-dependent treatment strength. The growth rate-independent treatment instead causes a cancer cell mortality rate *m* that is equal for all cells. It could resemble a type of immunotherapy that targets a surface protein that is present on all cancer cells. These assumptions result in the following system of differential equations describing the change in the sizes of the subpopulations,

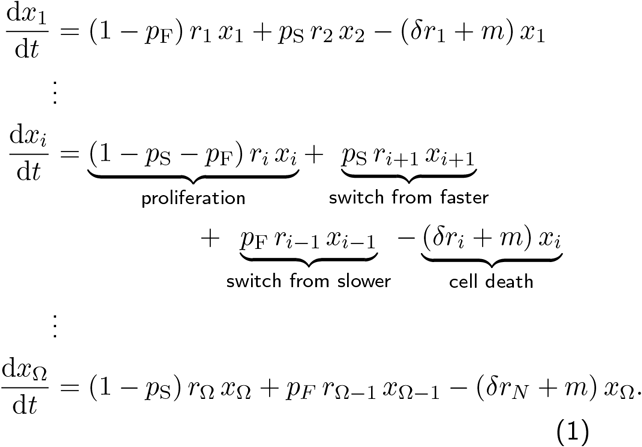

**Figure 1.**
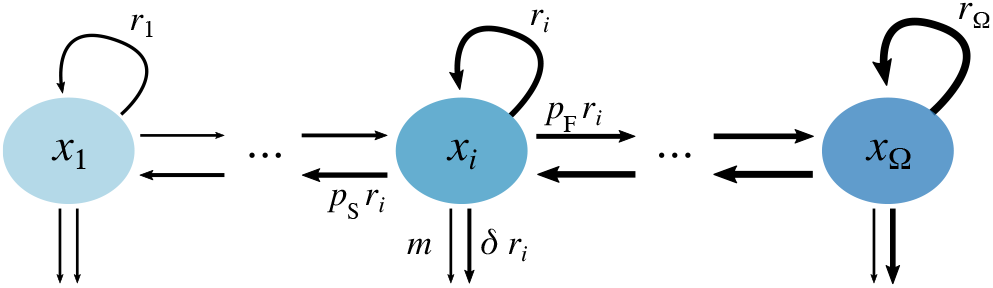
Model sketch. Arrows indicate growth, switching and death processes of the different subpopulations. Cells grow at rates *r_i_*, switch to slower growth rates at rate *p_S_ r_i_* or to faster growth rates at rate *p_F_ r_i_*. Cancer cell mortality from growth rate-dependent treatment (for example by the uptake of chemotherapeutics) is assumed to be proportional to growth rate and induces a mortality *δr_i_*. Cancer cell mortality from the growth rate-independent treatment, for example immunotherapy, is captured by the death rate *m*.

In our model, contributions of individual subpopulations to the whole population, and therefore also the resulting trait distribution, will converge to a stable distribution in time (S1). In principle, this distribution can be calculated analytically. Already for Ω > 2, however, the resulting expressions become unhandy and provide little further insight. For our purposes, it is sufficient to know that this stable distribution exists and is reached numerically.

We assume that the growing tumour will have approached this stable trait distribution before cancer diagnosis. After diagnosis, a period of treatment is applied. After treatment is halted, the cancer is monitored for an additional period to track the potential relapse dynamics. Before detection and after treatment termination, both the trait-independent and the trait-dependent treatment parameters (m and δ) are set to zero. Only during the treatment phase, one of them is set to the reference value from Tab. 1, depending on whether the trait-independent or the trait-dependent treatment is applied. We assume that both treatment types reduce the tumour load. To allow comparison of the two treatment types, we chose the cancer cell mortality rate from trait-independent treatment *m* such that the total cancer cell population after applying either of the two treatments is approximately equal at the end of the treatment phase. This ensures that both treatment types result in the same tumour load reduction, allowing a better comparison for our purposes. The slowest and fastest growth rates *r*_min_ and *r*_max_, as well as the switching parameters *p_S_* and *p_F_*, are chosen such that within the simulated treatment phase the slowest subpopulation can exceed the fastest subpopulation under trait-dependent treatment.

The system of differential equations (Eqs. 1) is numerically integrated for Ω = 25 subpopulations using the *LSODA* implementation of the *solve_ivp* function from the Scipy library [31] in Python (version 3.7). To equilibrate the ratios of adjacent subpopulations and arrive at the initial stable trait distribution, we first integrated for 200 time units from an exponential trait distribution 10^-60^e^80V_i_^, where *v* is an array of Ω linearly increasing values between 0 and 1. The result is taken as the initial condition for the pre-detection period.

### Treatment schemes

We investigate different predefined treatment schemes, where either only one treatment type is applied for the whole duration of treatment or the two treatment types are alternating. Additionally, we study an adaptive treatment scheme where at regular re-evaluation intervals Δ*t* the treatment type that induces the higher mortality on the total cancer cell population 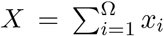 is chosen and continued until the next treatment re-evaluation. To make this decision we consider the rate of change of *X*

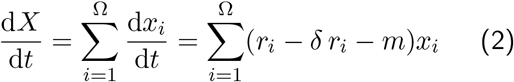

where the two last terms represent the mortality exerted by trait-dependent and trait-independent treatment, respectively. Which of these terms is larger depends on the trait distribution, which may change over time. For the adaptive treatment scheme, we evaluate those two terms and set the treatment type that exerts the lower mortality to zero. Maximum mortality is achieved by continuously re-evaluating the treatment type (Δ*t* → 0), which leads to the optimal adaptive treatment scheme.

### Stochastic model

During the treatment phase, the number of cancer cells typically drops drastically. As the cancer cell population is driven to low numbers, the population dynamics are affected by stochasticity and the mean-field approximation for large cell numbers becomes invalid. A deterministic model cannot capture true extinctions of the cancer population unless an extinction threshold is defined, which still fails to capture stochastic effects. To mechanistically capture this stochastic regime of low cancer cell numbers, we therefore develop a stochastic formulation in parallel to the deterministic model described above. Importantly, the deterministic and the stochastic model are based on the same microscopic processes and therefore directly comparable. However, for the stochastic model we constrain ourselves to only the two extreme subpopulations that grow at rates *r*_min_ and *r*_max_, as their dynamics will show the most pronounced differences. To obtain the stochastic trajectories, we simulate the microscopic processes stochastically using the Gillespie algorithm implementation in StochKit [32] for 10^4^ replicate populations. Additionally, we solve the stochastic differential equivalent to Eq. 1 numerically using the *sdeint* package (Matthew J. Aburn, version 0.2.1). For the derivation of the stochastic model, we refer to the Supplementary Material. All computational implementations can be found at [DOI: 10.5281/zenodo.4461667]. The data is available at [DOI: 10.5281/zenodo.4293320].

## Results

We represent a tumour as a population with a range of different growth rates. This allows us to infer how growth rate-dependent treatment affects population decline and relapse dynamics differently from growth rate-independent treatment. We find that the differential effect of the growth rate-dependent treatment changes the relative abundances of the subpopulations (Fig. 2a) and changes the trait diversity within the population resulting in two diversity peaks: one during treatment and one during early relapse (Fig. 2b). This is different from the trait-independent treatment, where diversity is constant. Before detection, fast growth rates are selected for and dominate the population at detection (Fig. 2, S1). Growth rate-dependent treatment, while inducing a decline in total cancer cell numbers, selects for slower growth rates which eventually allows the slowest growing subpopulation to take over the population. The timing of this take-over depends on the stable trait-distribution, which is determined by the rates of switching along the trait axis due to phenotypic plasticity or genotypic variability (Supplementary Material). After treatment termination, all subpopulations resume to grow at their respective growth rates. Thus, faster subpopulations quickly take over the population again (Fig. 2). Cell switching creates a net influx from faster to slower sub-populations. Thus, also the slow-growing subpopulations eventually increase at almost the maximum growth rate (Fig. 2, S2). However, the subpopulations with the fastest growth rates outnumber the slower cells by orders of magnitude (S1). For growth rate-independent treatment, the high relative abundance of fast-growing cells is not affected by treatment.

**Figure 2.**
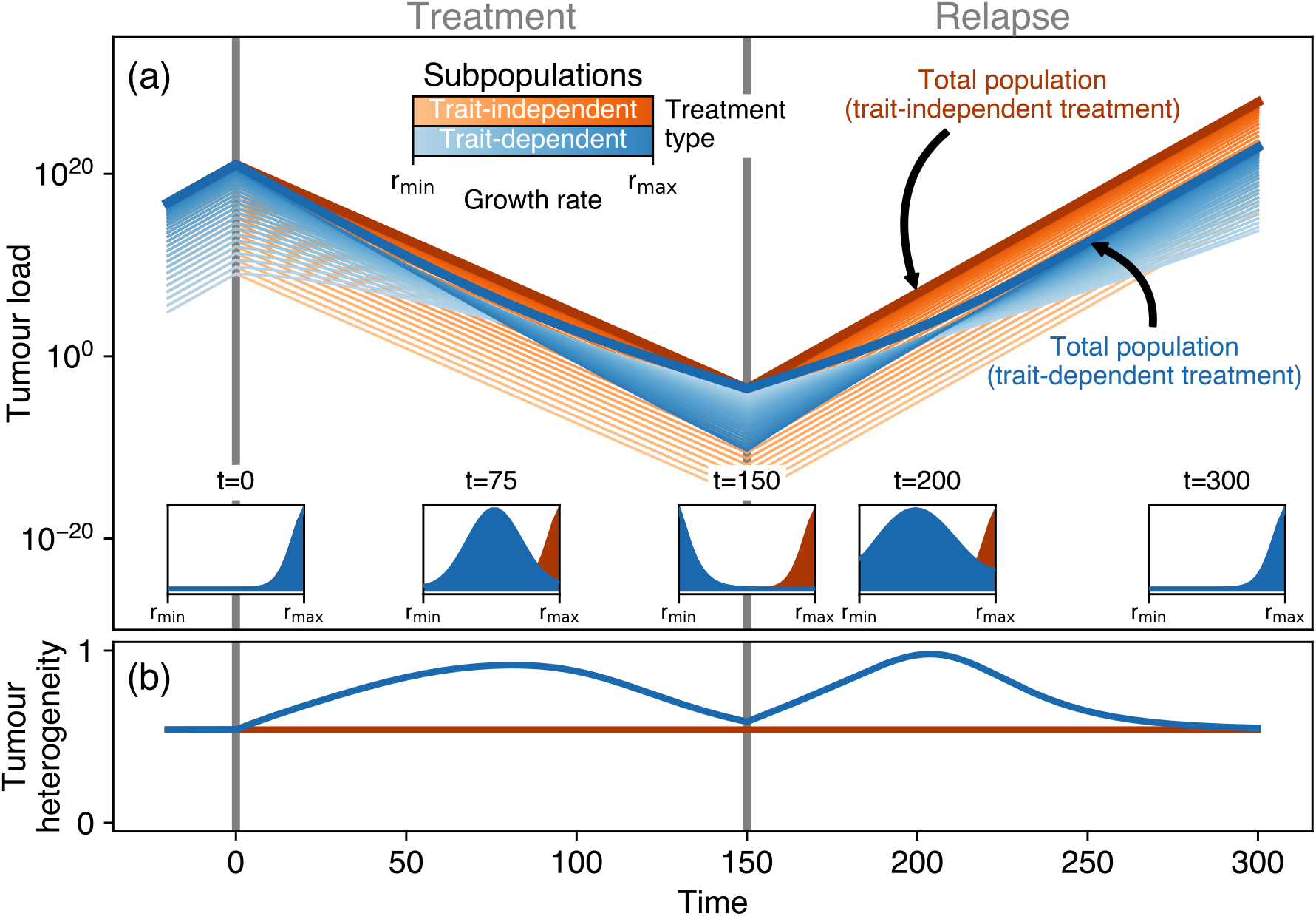
Population dynamics for a typical treatment scenario. Starting from a cancer population that reached the stable distribution before detection at *t* = 0 treatment is applied between *t* = 0 and *t* = 150. After *t* = 150 relapse is monitored until *t* = 300. (a) Splitting the total cancer population (thick lines) into different subpopulations (thin lines) with different growth rates (colour gradient) allows for tracking the differential selection pressure that trait-dependent and trait-independent treatment types impose on different growth rates and how this selection affects the subsequent relapse dynamics. Insets show the growth rate trait distribution at various time points. The cancer cell mortality rate in the trait-independent treatment was set such that the tumour load at the end of treatment is similar to the tumour load at the end of the trait-dependent treatment. (b) Trait diversity (measured as Shannon evenness) is affected only by the growth rate-dependent treatment.

**Table 1.** Reference parameter set. Deviations from these values are reported where applicable.

| Parameter | Biological meaning | Value |
| --- | --- | --- |
| $r_1 = r_{\min}$ | Growth rate of the slowest subpopulation | 0.25 time unit<sup>-1</sup> |
| $r_{\Omega} = r_{\max}$ | Growth rate of the fastest subpopulation | 0.5 time unit<sup>-1</sup> |
| $p_S$ | Factor scaling the switching to slower subpopulations | 0.2 |
| $p_F$ | Factor scaling the switching to faster subpopulations | 0.2 |
| $\delta$ | Cancer cell mortality factor of the trait-dependent treatment | 2 |
| $m$ | Cancer cell mortality rate of the trait-independent treatment | 0.86 time unit<sup>-1</sup> |

The growth rate of the most abundant subpopulation sets the speed of relapse. Accordingly, we observe a biphasic relapse pattern after the termination of the trait-dependent treatment (Fig. 2a). As long as the slowest subpopulation remains most abundant, the total cancer population grows at a slow rate, but as soon as the fast subpopulation takes over, also the whole population increases at the maximum growth rate (S2). This particular relapse behaviour contrasts with the relapse pattern for a trait-independent treatment. Assuming a comparable treatment effect, i.e. treatment reduces the total tumour load by the same amount, we see that here, the fastest subpopulation, although declining, remains dominant throughout treatment and during relapse. Thus, also the relapse occurs at the fastest growth rate immediately after treatment termination for trait-independent treatment types. For comparable tumour load reductions, our model therefore predicts that a potential relapse after growth rate-dependent treatment occurs substantially later than after growth rate-independent treatment.

Such relapse, however, is subject to stochasticity. Towards the end of our simulated treatments, cancer cell numbers become low and stochastic extinction of subpopulations, as well as the eradication of the whole cancer population, can occur. To illustrate this behaviour, we conducted stochastic simulations for a simplified model with only two subpopulations, comparing growth rate-dependent and growth rate-independent treatment in a large number of replicates. Again, we ensured that the total reduction of tumour cells in both treatments is the same. We find that for the trait-independent treatment type, the tumour goes extinct in more replicates (Fig. 3), while for the trait-dependent treatment type the slow subpopulations quickly take over and prevent extinctions in many cases. After the treatment is terminated, the extinct fast subpopulations of the surviving replicates first need to be repopulated from the slow subpopulation, leading to a further delay of relapses. Only then the tumour regrows at the speed found in the deterministic model. Accordingly, relapse on average occurs later for the trait-dependent treatment type in the stochastic simulations, in agreement with the findings from the deterministic model (Fig. 4). For the extreme case of no switching between subpopulations, relapse would proceed only at the growth rate of the slow subpopulation. If the switching rate is very low, it can take a considerable amount of time until relapse proceeds at the rate of the fast subpopulation again. In the deterministic description, however, the increase of the fast subpopulation is only speeded up by, but not contingent on, the switching of cells from the slow into the fast subpopulation. Thus, relapse inevitably proceeds at the growth rate of the fast subpopulation eventually. Relapse will thus always show a biphasic pattern in a deterministic description, but it might not in a stochastic description or when the mean-field approximation is invalid.

**Figure 3.**
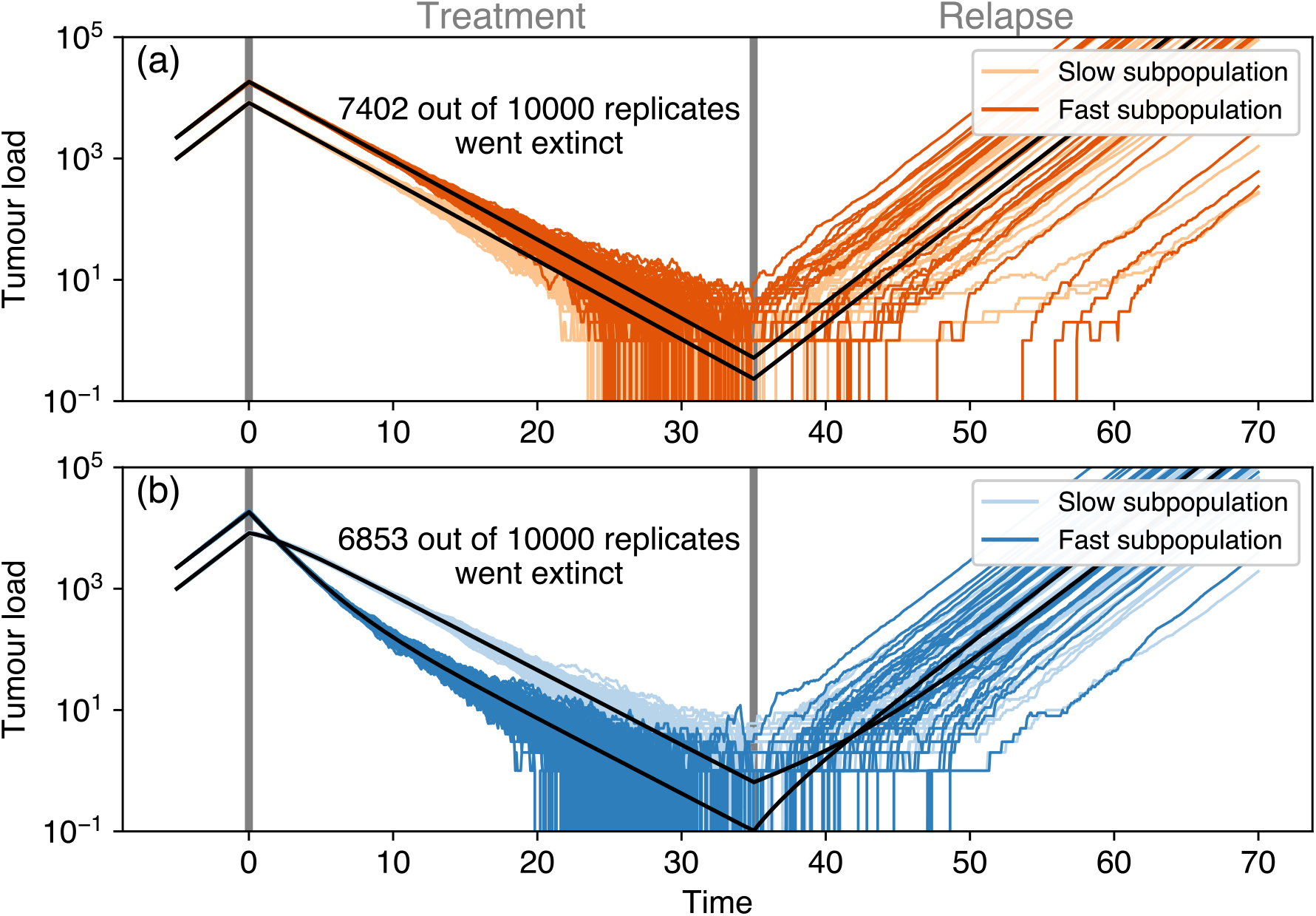
Stochastic simulations. Shown are 100 replicate populations for a binary trait where the two subpopulations grow with growth rates *r*_min_ and *r*_max_, respectively, for (a) trait-independent and (b) trait-dependent treatments. The black lines represent the deterministic solution of the ordinary differential equation for the two subpopulations (Eq. 1). Note that initial conditions and treatment duration are different compared to Fig. 2 to allow for relapse given the discrete number of cells in the stochastic simulations. The cancer cell mortality by trait-independent treatment was set to *m* = 0.722 d^-1^ to ensure equal tumour load at the end of treatment in the deterministic model for the shorter treatment duration.

**Figure 4.**
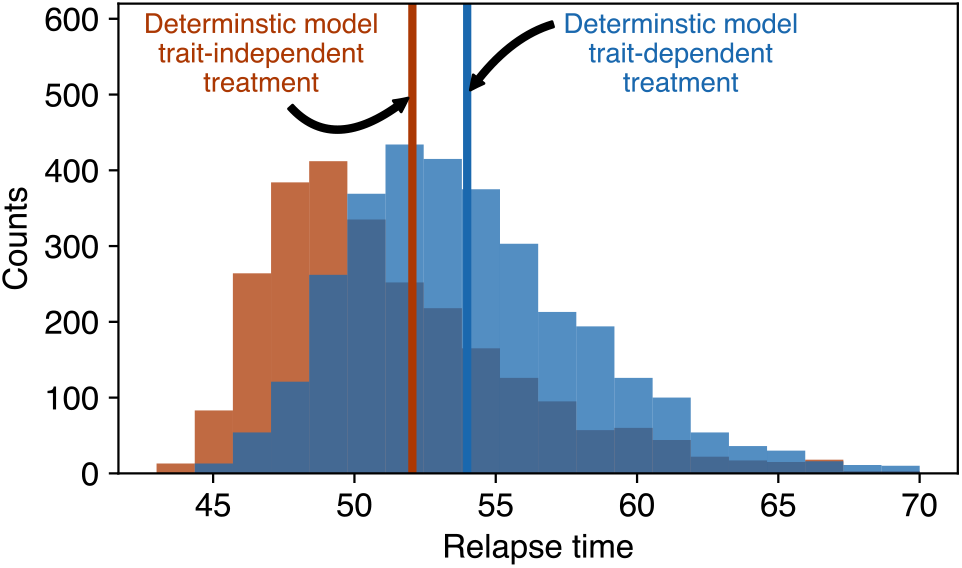
Relapse time distributions extracted from the stochastic simulations in Fig. 3. Orange represents the trait-independent and blue the trait-dependent treatment type. The vertical lines indicate the relapse times from the deterministic simulations. Relapse is defined to occur when the total tumour load of a replicate exceeds 10^3^ cells.

To investigate how the stochastic contributions from the slow and fast subpopulations differ, we integrated only the stochastic term in the stochastic differential equation (Eq. S1.6) while setting the deterministic term to zero. We find that the slow subpopulation explores a smaller state space range by taking smaller steps (S3). The slow subpopulation may therefore act as a refuge against extinction as it would require more time to eventually cross the extinction boundary where the cancer cell number drops to zero. As growth rate-dependent treatment increases the diagonal entries of the diffusion matrix in Equation S1.6 proportionally to the growth rates, it also increases stochastic step sizes proportionally to the respective growth rates. Therefore, under growth rate-dependent treatment, the steps that the fast subpopulation is taking will be even larger than the steps of the slow subpopulation, making the extinction of the fast subpopulation even more likely than the extinction of the slow subpopulation. Growing slowly thus reduces a cell’s chance to be killed by growth rate-dependent treatment. For the trait-independent treatment, however, we find that the differences between the stochastic step sizes of the slow and fast subpopulations become smaller compared to no treatment (S3). Here, the treatment-induced mortality is equal for both subpopulations and dominates the diagonal entries of the diffusion matrix in Equation S1.6. This decreases the relative differences between the stochastic step sizes of both subpopulations, which undermines the refuge effect of the slow subpopulation.

So far, we have assumed that only a single treatment type may be chosen for the whole treatment duration. Even if toxicity or inhibiting interactive effects may prevent simultaneous application of a trait-dependent and a trait-independent treatment, their sequential application is often feasible. We find that by appropriately choosing the treatment sequences, increased chances of cure and delay of relapse may be achieved (Fig. 5). To increase the chance of cure, a treatment sequence should be chosen that maximizes cancer cell mortality. To delay relapse, the trait distribution should be maximally shifted towards slow growth rates. This results in two different treatment goals, which can only partly be met by the same treatment scheme.

**Figure 5.**
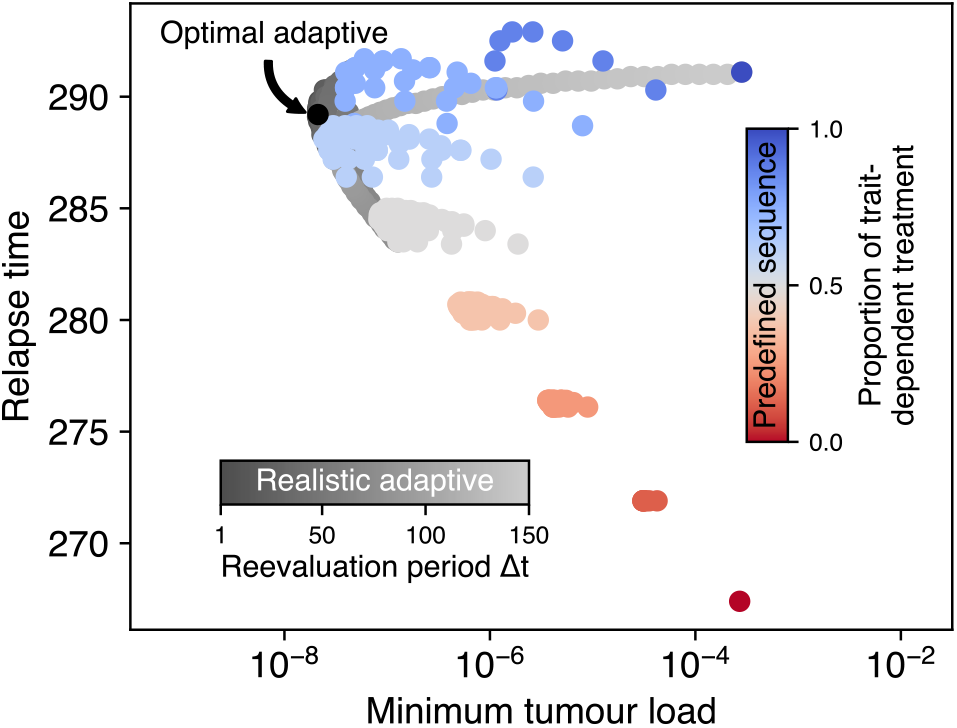
Comparison of minimum tumour load during treatment and the relapse time when tumour load surpasses the pre-treatment maximum for different sequential treatment schemes. Alterations between the trait-independent and the trait-dependent treatment type are fixed in the predefined sequence scheme (red to blue colour gradient corresponds to higher proportion of trait-dependent treatment type). In the realistic adaptive scheme, the currently best treatment type is determined at regular intervals during the treatment phase (grey colour gradient). In the optimal adaptive scheme (black dot), the treatment type that, given the current trait distribution, would exert the highest population mortality is chosen nearly instantaneously (at every step of the numerical solver). Note that these schemes have a much stronger impact on the minimal tumour load (up to a factor of 1000) than on the relapse time (up to a factor of 1.3).

We studied both predefined sequential treatment schemes where trait-independent and traitdependent treatment alternate (S4, S5) as well as adaptive schemes where the trait distribution within the tumour is re-assessed at regular intervals *Δt* (realistic adaptive schemes, S6). Following the assessment, the treatment is continued with the treatment type that maximizes the mortality of the cancer cell population given the current trait distribution (S7). Additionally, we include an optimal adaptive scheme that employs trait distribution assessments at a very high frequency Δ*t* → 0 as an extreme case.

We find that the optimal adaptive scheme indeed minimizes the tumour load at the end of the treatment phase. However, the predefined and realistic adaptive schemes can achieve a slightly longer time to relapse (Fig. 5). Interestingly, we find that already intermediate reevaluation periods in the realistic adaptive scheme and even some predefined sequences result in treatment results close to the theoretical optimum. For the predefined scheme, latest relapse is achieved by including a short period of trait-independent treatment in the middle of an otherwise trait-dependent treatment (S4). Lower minimum tumour loads are achieved by a combination of frequent treatment switching and a higher proportion of trait-dependent treatment. The adaptive scheme with realistic reevaluation periods Δ*t* generally approaches the optimal adaptive scheme for Δ*t* → 0 and converges to the pure trait-dependent scheme as the reevaluation period becomes large (S6). At intermediate Δ*t* we observe multiple peaks in both the minimum tumour load and relapse time wherever the total treatment duration is an integer multiple of Δ*t* and the number of possible switches changes. For example if the treatment reevaluation period is between half of the total treatment duration and the total treatment duration, then only a single switch of treatment type is possible. In contrast, for only slightly smaller reevaluation periods two switches are possible. If there are only few switches, they can have strong effects on the trait distribution and thus give rise to discontinuities in the minimum tumour load and relapse time. At the onset of treatment, the trait distribution is heavily skewed towards large growth rates (S7). The optimal and all adaptive sequential schemes initially apply the trait-dependent treatment, thus driving the mean of the trait distribution to intermediate values, as already observed in Fig. 2. This eventually decreases the population mortality rate from trait-dependent treatment. Accordingly, the optimal treatment sequence switches to the trait-independent treatment when the population mortality rates for both treatment types become equal (Eq. 2), which implies

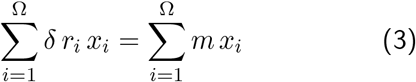

As now the trait distribution is freed from trait-dependent selection, the mean of the trait distribution increases as the faster-growing subpopulations increase in frequency, which eventually favours the trait-dependent treatment again. By continued rapid switching of treatment types, the optimal adaptive scheme modulates the trait distribution and maintains the mean trait value of maximum cancer cell population mortality 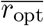, which follows from Eq. 3 as

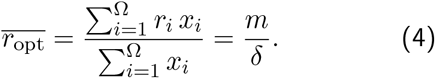

The mean trait value for the realistic adaptive treatment scheme fluctuates around 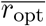 and approaches it for small Δ*t* (S7).

## Discussion

Understanding cancer as a population of phenotypically diverse cells suggests representing this population by a trait distribution of considerable variance, on which treatment types can select. In this study, we have entirely focussed on growth rate as the focal trait. We compared a growth rate-dependent and a growth rate-independent treatment. The growth rate-dependent treatment is motivated by chemotherapy: Many chemotherapeutic drugs are cell-cycle specific and only damage dividing cells. Accordingly, we hypothesized that chemotherapy targets fast proliferating cells preferentially and exerts higher mortality on them. Our growth rate-independent treatment is motivated by immunotherapy: This therapy targets cancer cells irrespective of their proliferation rate, for example by using bi-specific antibodies that specifically label cancer cells, which are then recognized and killed by the immune system. Slow and fast cells are therefore equally targeted by the immune system. Even though these hypotheses are likely to hold in many cases, they might not generally apply, but depend on specifics of the particular cancer, treatment types and patient.

In the case of acute lymphoblastic leukaemia, where tracking the proportion of malignant cells over time is possible, it was found that chemotherapy often leaves behind a small number of malignant cells, a situation termed minimal residual disease. The presence of this minimal residual disease is of high prognostic value and indicates a high likelihood of future relapse [29, 33, 34]. In such cases, it was found that Blinatumomab, a bi-specific monoclonal antibody, can often suppress this residual disease below detection levels [35]. For patients with relapsed or refractory B-cell precursor acute lymphoblastic leukaemia that already underwent multiple chemotherapy treatments, switching to immunotherapy with Blinatumomab showed significantly better treatment outcomes than conducting additional chemotherapy [36]. This seems reasonable under our assumption that chemotherapy would shift the growth rate trait distribution to smaller values, where additional chemotherapy only has limited effect. Choosing a different treatment type that does not select for the same trait would allow for a further and stronger reduction of tumour load. The relapsed/refractory setting thus resembles one of the close-to-optimal treatment schemes where towards the end of the treatment phase a period of growth rate-independent treatment is introduced, after prior growth rate-dependent treatment has shifted the trait distribution to values of decreased sensitivity against the trait-dependent treatment. Front-line approaches of using combinations of chemotherapy and immunotherapy are also promising and show improved treatment effects compared to chemotherapy alone [37–40]. Complementing reports of overall survival data with time series of malignant cell counts, as for example in [41], could provide mechanistic insights into why and how these combination therapies work. Phenotypic trait distributions could be different after chemotherapy and immunotherapy, despite resulting in the same minimal residual disease. This may contribute to an explanation of why the prognostic value of minimal residual disease levels could be different for these two treatment alternatives.

We have found that slower proliferating subpopulations may present a refuge during chemotherapy, from which relapse may arise. While accounting for the full trait distribution is essential to understand this pattern, detecting it requires only knowledge about the time course of the total tumour load. The fingerprint for this scenario of slow populations being sheltered from treatment are the biphasic dynamics (or multiphasic) of tumour load both during trait-dependent therapy and relapse [42, 43]. During treatment, the initial tumour load decrease is driven by the effective growth rate of the fastest-growing subpopulation. In contrast, the effective growth rate of the slowest-growing subpopulation determines the rate of tumour load decrease towards the end of treatment. The situation inverts during relapse with the slowest growing subpopulation setting the rate of increase initially before finally the effective growth rate of the fastest growing subpopulation determines the speed of relapse. Advances in sampling precision and frequency will eventually provide a temporal resolution of the total tumour load also in clinical settings. This may also allow the detection of biphasic (or multiphasic) dynamics, which could act as the fingerprint for phenotypic heterogeneity among cancer cells and guide appropriate treatment decisions, notably only requiring total tumour load, not the trait distribution itself.

If such a pattern of changing dominance would be detected, our results predict that switching to a different treatment type that does not select on the same trait as the previous treatment will improve treatment effect by allowing stronger tumour load reduction and delayed relapse. Interestingly, we have seen that also larger and more realistic check-up intervals would suffice for close-to-optimal treatment effects, a finding that was also observed for other adaptive treatment schemes, such as tumour containment [25]. Since longer check-up intervals would lead to a substantial growth above the clinical detection limit the precise value of this detection limit is not a crucial determinant for the success of the adaptive scheme.

On a more abstract level, however, we have combined two treatment types, one independent of, the other dependent on a certain characteristic (the focal trait) of the cancer cells, with the trait-dependent type offering a route for resistance. This creates an evolutionary double bind by the two treatment types as the resistance mechanism of decreasing growth rate is countered by relaxing the trait-selective treatment [17]. Then, due to their higher growth rate, faster-growing subpopulations will increase again, which automatically restores sensitivity. This alone would correspond to the adaptive treatment approach [20]. Filling the treatment break with a second, trait-independent treatment does not hinder the favourable overtake by the more susceptible faster growing subpopulation and further decreases the tumour cell numbers. Building on the established idea of targeting specific phenotypes in cancer treatment [8, 44] and the notion of the prevalence of intratumour heterogeneity, our approach shows how to tailor personalized treatments to the phenotypic trait distribution of cancer cells.

## Acknowledgements

We thank Weini Huang and Benjamin Werner for their feedback during the initial phases of this project and Luka Opašić for helpful comments on an earlier version of this manuscript.

## Supplementary Information

## S1 Appendix Derivation of the stochastic model

Here we consider a stochastic model of tumour growth with two phenotypes, namely, slow and fast proliferating tumour cells. These subpopulations grow and switch their phenotypes at different rates. When the trait-dependent treatment is applied, they die at rates proportional to their growth rate. The trait-independent treatment induces the same mortality rate on both phenotypes. We start the derivation of the stochastic model by presenting these processes as a set of reactions that individual cells perform. The reactions are accompanied by corresponding reaction rates *ρ_k_* and the change vector ***μ***_k_, that describes the effect of a single reaction. The reactions can be implemented in a stochastic simulation algorithm [1].

**Table 2:**
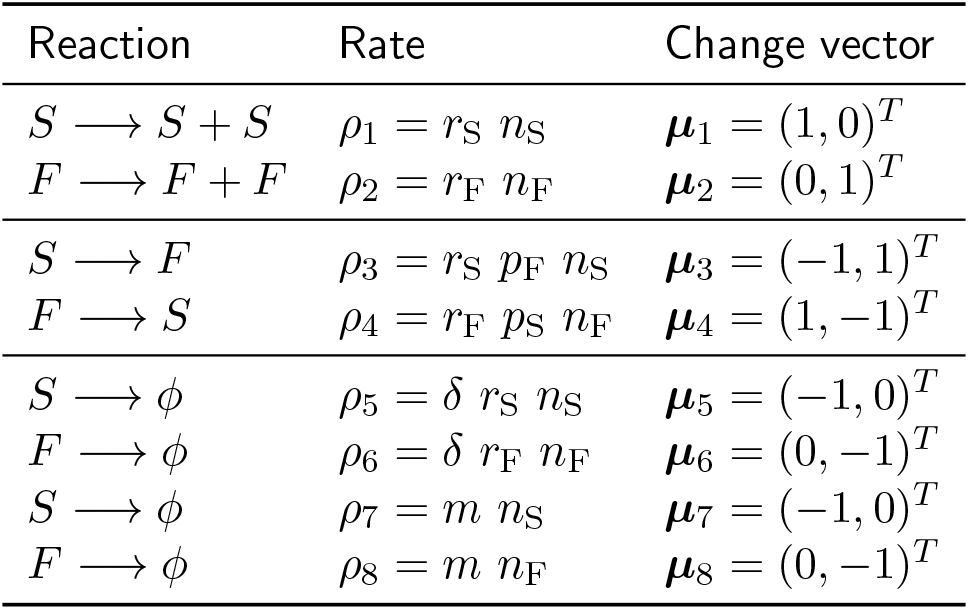
The eight possible reactions in the stochastic model. Reactions going to *φ* represent cell death events. *S* and *F* represent individual cells, *n_S_* and *n_F_* are their total numbers in the cancer cell population. Reaction *k* occurs with rate *ρ_k_* and changes the cell number vector ***n*** = (*n_s_,n_p_*) by the change vector ***μ***_k_.

| Reaction | Rate | Change vector |
| --- | --- | --- |
| $S \longrightarrow S + S$ | $\rho_1 = r_S n_S$ | $\mu_1 = (1, 0)^T$ |
| $F \longrightarrow F + F$ | $\rho_2 = r_F n_F$ | $\mu_2 = (0, 1)^T$ |
| $S \longrightarrow F$ | $\rho_3 = r_S p_F n_S$ | $\mu_3 = (-1, 1)^T$ |
| $F \longrightarrow S$ | $\rho_4 = r_F p_S n_F$ | $\mu_4 = (1, -1)^T$ |
| $S \longrightarrow \phi$ | $\rho_5 = \delta r_S n_S$ | $\mu_5 = (-1, 0)^T$ |
| $F \longrightarrow \phi$ | $\rho_6 = \delta r_F n_F$ | $\mu_6 = (0, -1)^T$ |
| $S \longrightarrow \phi$ | $\rho_7 = m n_S$ | $\mu_7 = (-1, 0)^T$ |
| $F \longrightarrow \phi$ | $\rho_8 = m n_F$ | $\mu_8 = (0, -1)^T$ |

As this infinitesimal time element tends to zero the discrete model leads to a stochastic differential equation model [2, Section 5.1]. To get there, we compute the vector of expected change **μ** to the population vector **n** per time unit as the sum over all possible change vectors weighted by their respective rates,

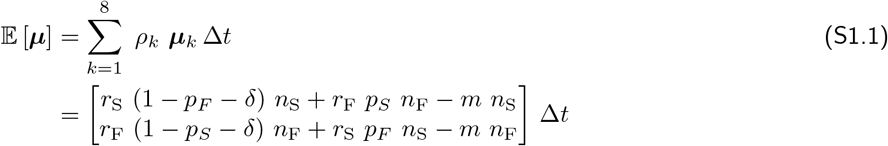

Similarly, we can obtain the covariance matrix of change rate to the population vector

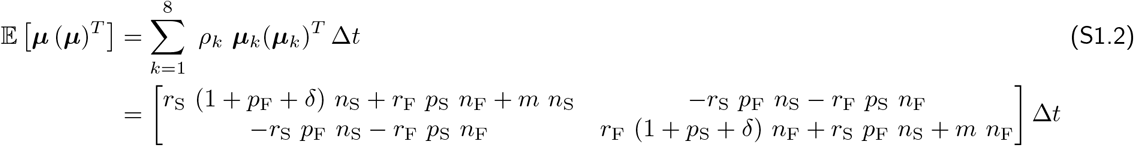

If we now introduce a system size parameter V and convert the numbers of cells ***n*** to densities 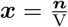, we can write the expectation vector *a*(*x*), and the covariance matrix ***B***(*x*) for the cell densities [2, Section 5.1],

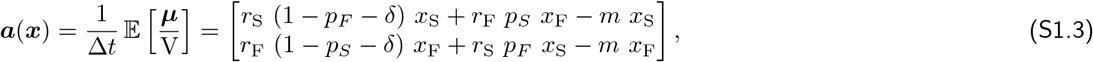

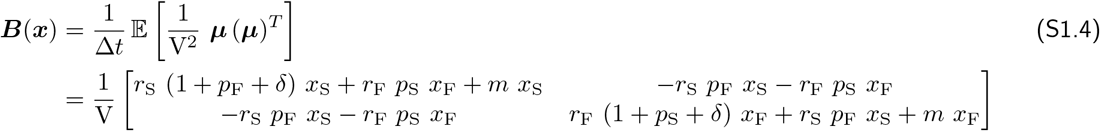

We arrive at an approximate Fokker-Planck equation that can be written as

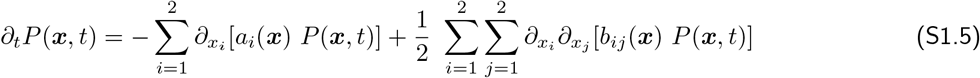

with *a_i_*(***x***) and *b_ij_*(***x***) being the entries of ***a***(***x***) and ***B***(***x***). We see that the system size parameter V determines the relative contributions of the drift and diffusion terms in the Fokker-Planck equation, thus setting the relative effect of stochastic fluctuations on the system dynamics. Using the Feynmann-Kac Formula, we finally obtain a system of Itô stochastic differential equations from the Fokker-Planck equation,

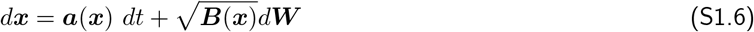

where ***W*** = (*W*_1_, *W*_2_)^*T*^ consists of two independent Wiener processes *W_i_*, or equivalently

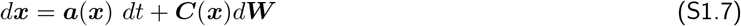

with ***C***(***x***)^*T*^***C***(***x***) = ***B***(***x***) [2, pp. 144]. This stochastic differential equation is numerically solved using the *sdeint* package (Matthew J. Aburn, version 0.2.1).

## S2 Appendix Effect of the switching parameters

The switching parameters *p_S_* and *p_F_* determine the flux between adjacent subpopulations. Smaller values broaden the stable trait distribution and prolong the time until both treatment types have achieved the same tumour load reduction, as it takes longer for the slowest subpopulation to overtake the fastest subpopulation (Fig. S2.1).

**Figure S2.1.**
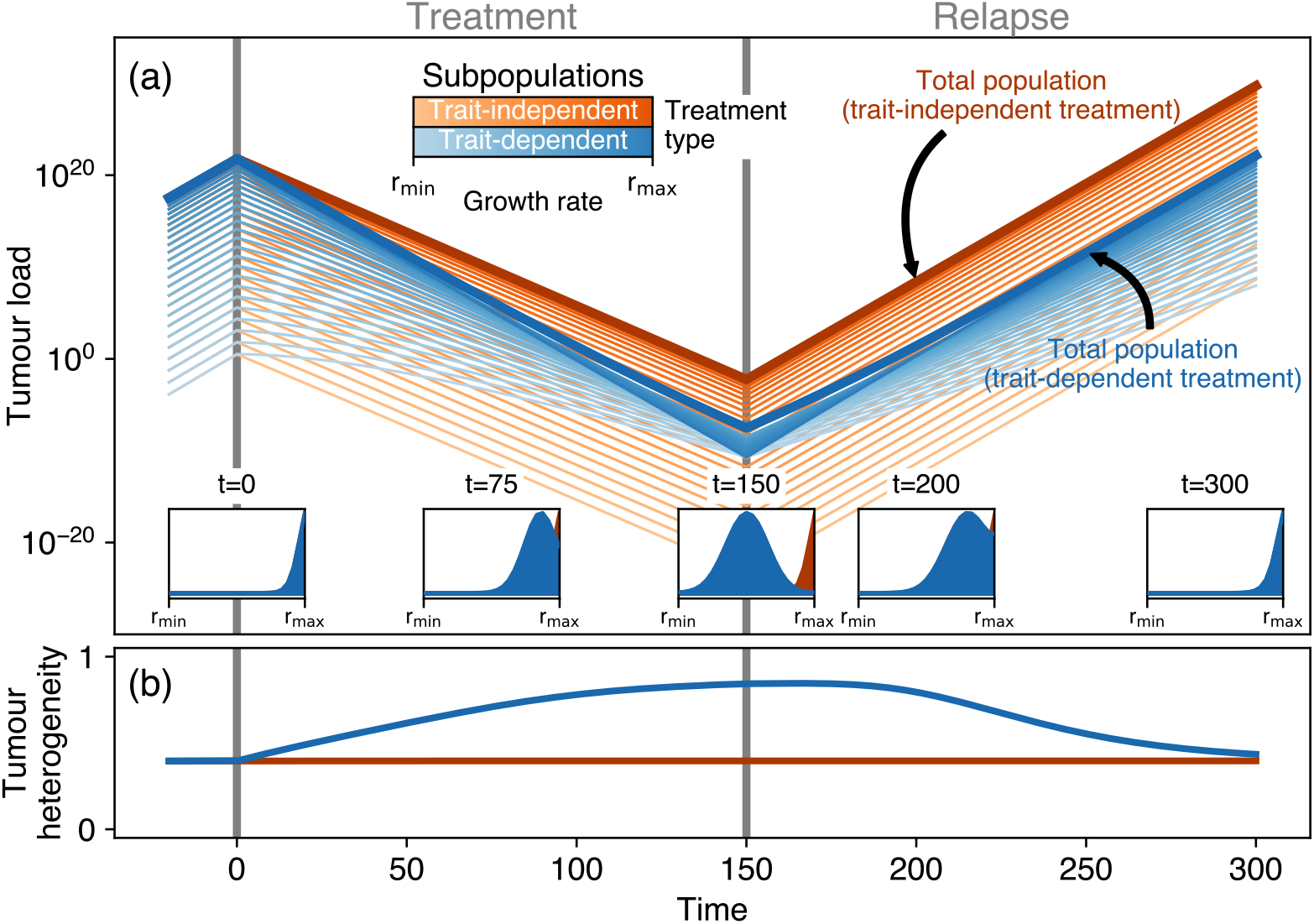
Same as Fig. 2 but for slower switching between adjacent subpopulations, *p_S_* = *p_F_* = 0.05.

If the switching parameters are larger, the stable trait distribution is narrower, and both treatment types achieve equal tumour load reductions earlier (Fig. S2.2). Throughout the paper, we have assumed that switching to adjacent subpopulations is equally likely. Asymmetric switching alters the stable trait distribution. If switching to faster-growing subpopulations is more likely than switching to slower-growing subpopulations, *p_F_* > *p_S_*, the trait distribution increases steeper towards faster growth rates (Fig. S2.3). As subpopulations with larger growth rates are now more populated, also the growth rate-dependent treatment has a higher effect than for symmetric switching. Also, the realized growth rate of the fastest subpopulation is higher, which increases the tumour load and decreases the effect of the trait-independent treatment type.

If switching to slower-growing subpopulations is more likely, the maximum of the trait distribution moves to slightly slower growing subpopulations, which decreases the realized growth rates and leads to lower tumour loads and therefore more effective treatment (Fig. S2.4).

**Figure S2.2.**
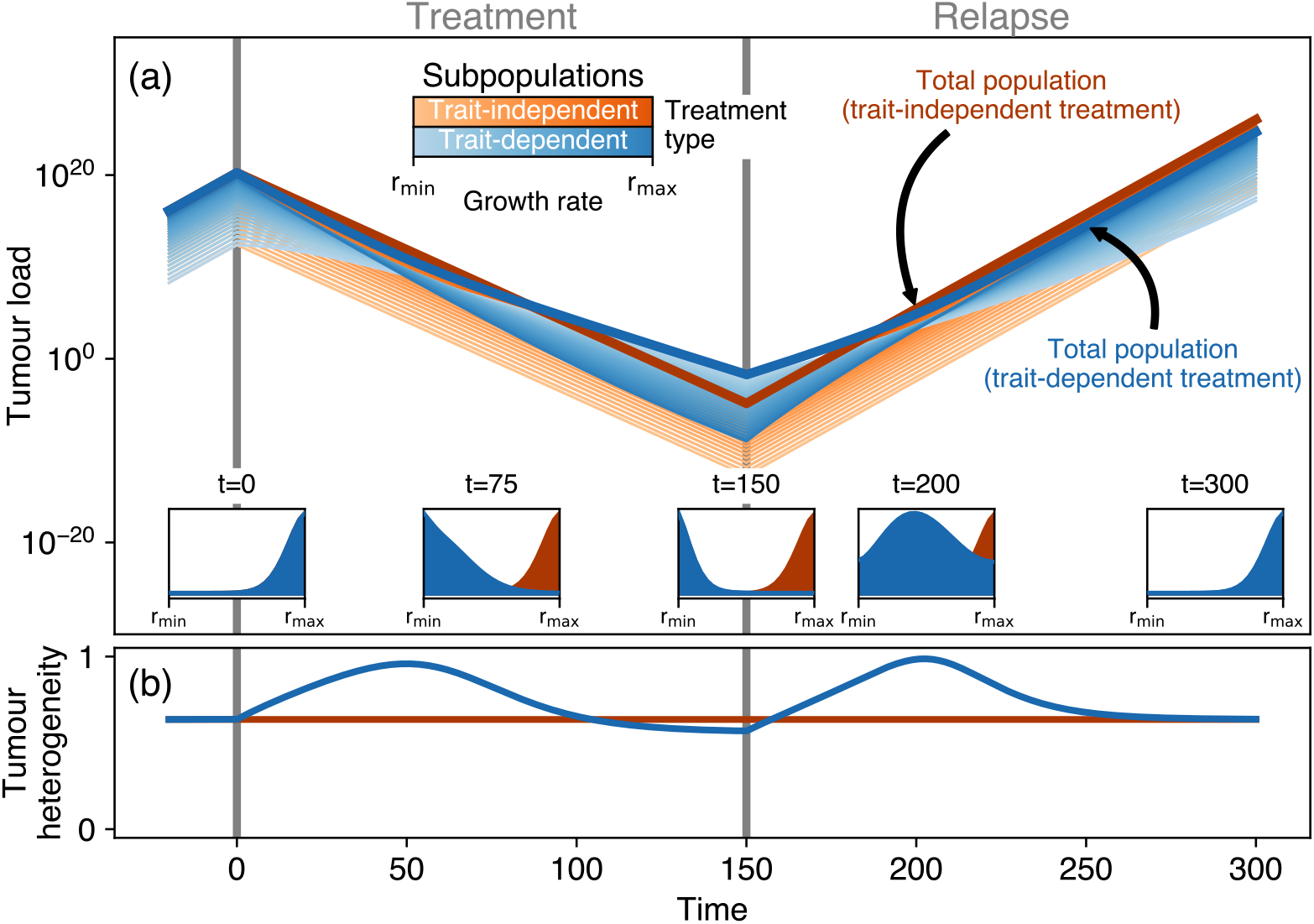
Same as Fig. 2 but for faster switching between adjacent subpopulations, *p_S_* = *p_F_* = 0.5.

**Figure S2.3.**
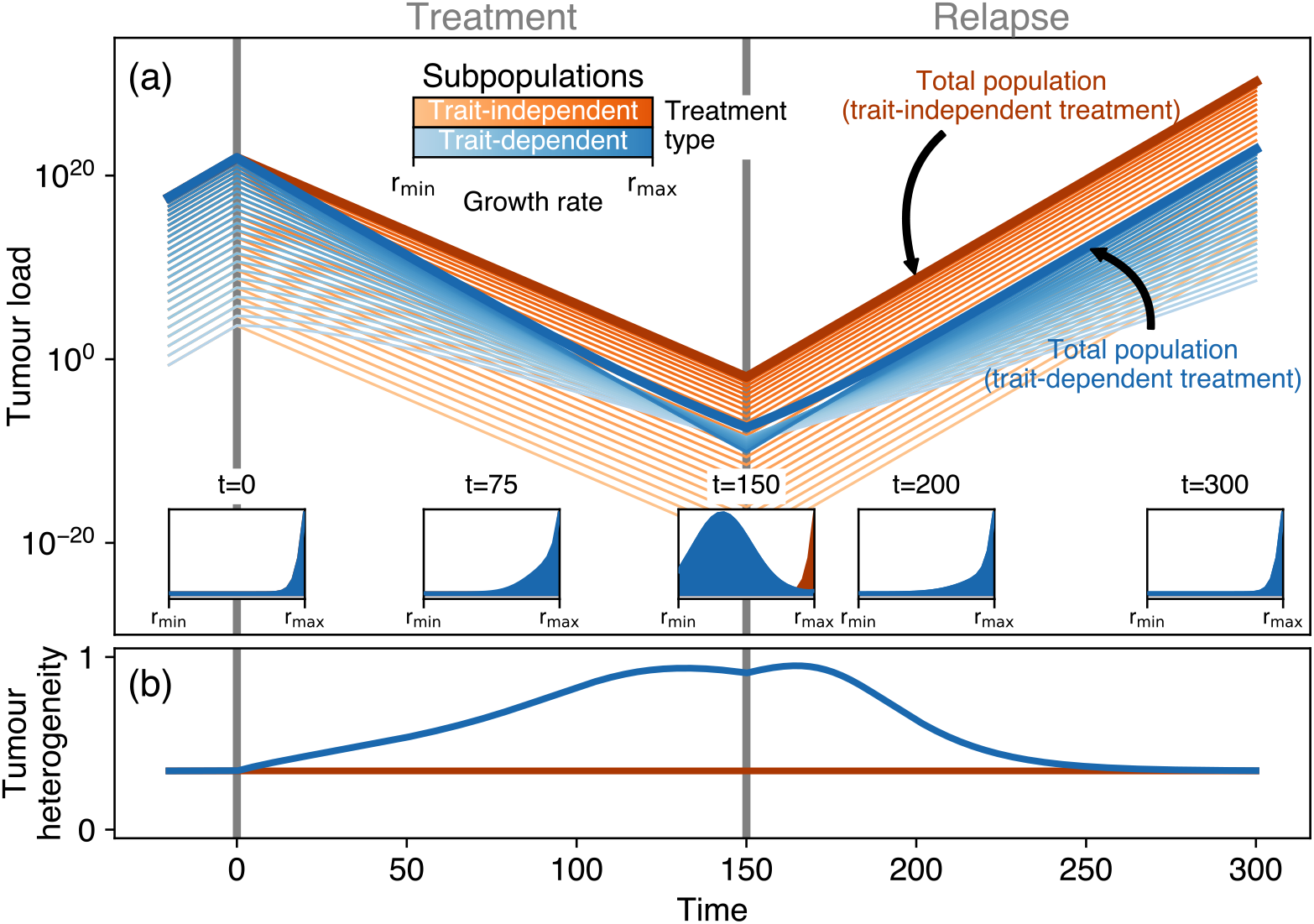
Same as Fig. 2 but now for asymmetric switching between adjacent subpopulations, assuming an increased flux to faster-growing subpopulations, *p_S_* = 0.1 and *p_F_* = 0.2.

**Figure S2.4.**
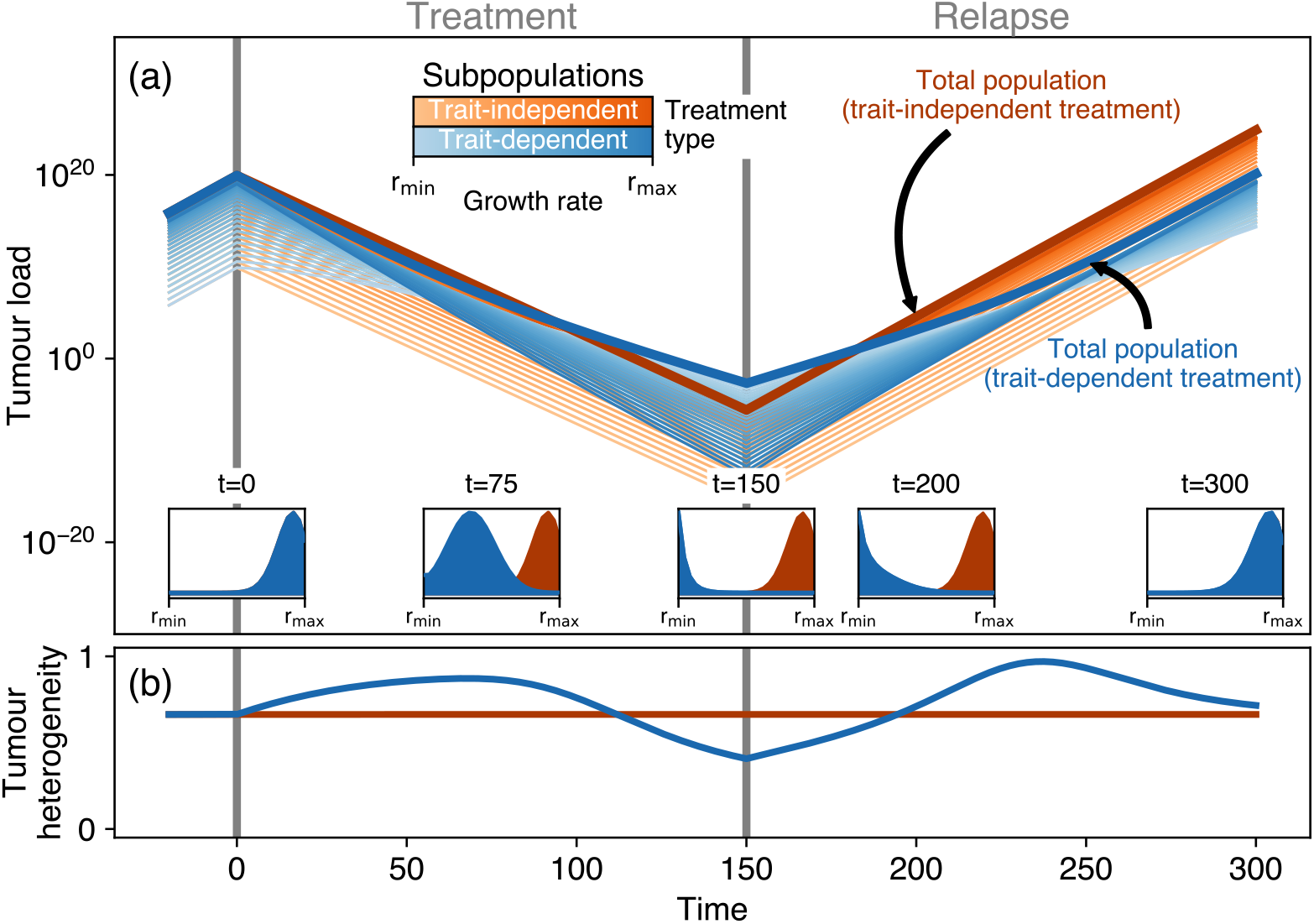
Same as Fig. 2 but now for asymmetric switching between adjacent subpopulations, assuming a decreased flux to faster-growing subpopulations, *p_S_* = 0.2 and *p_F_* = 0.1.

---

**Figure S1.**
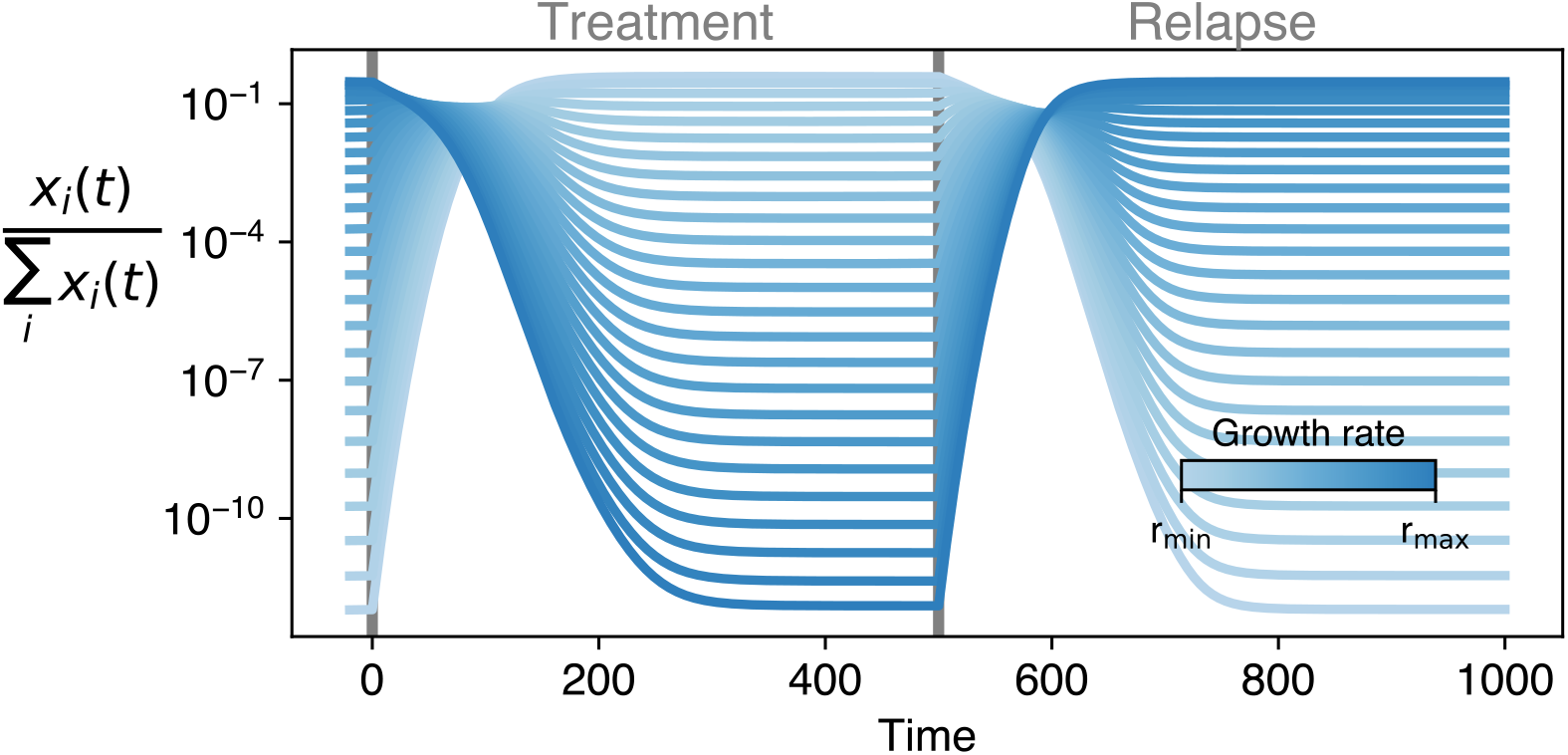
Relative contribution of every subpopulation for the trait-dependent treatment. Our model gives rise to a stable trait distribution (see constant ratios prior to the treatment phase). The trait-dependent treatment type creates another stable trait distribution towards the end of the treatment phase, where the slowest-growing subpopulations dominate. Note that treatment phase and relapse phase are prolonged here compared to Fig. 2 for better visualization.

**Figure S2.**
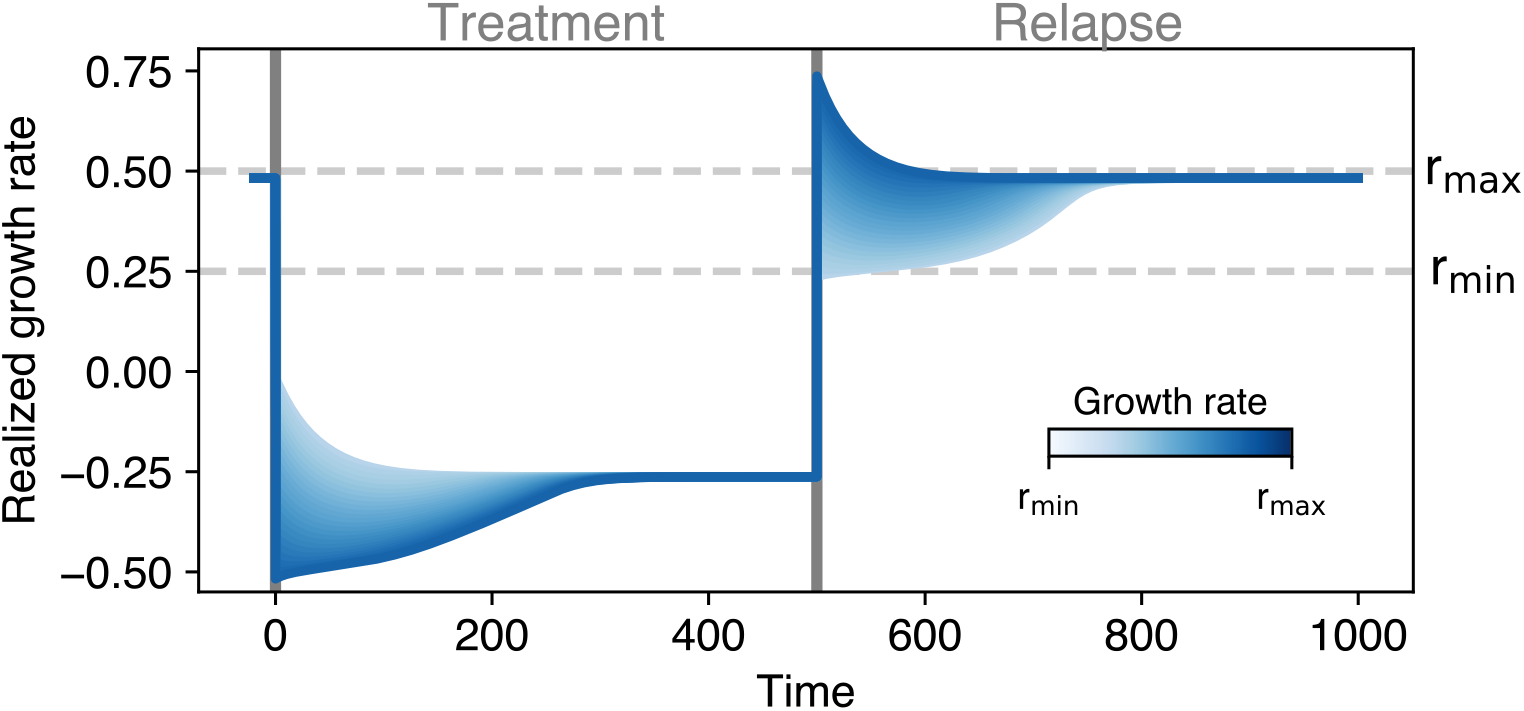
Realized subpopulation growth rates for the trait-dependent treatment. Switching to the adjacent slower subpopulation limits the realized growth rate of the fastest subpopulation to slightly below *r*_max_. Note that treatment phase and relapse phase are prolonged here compared to Fig. 2 for better visualization.

**Figure S3.**
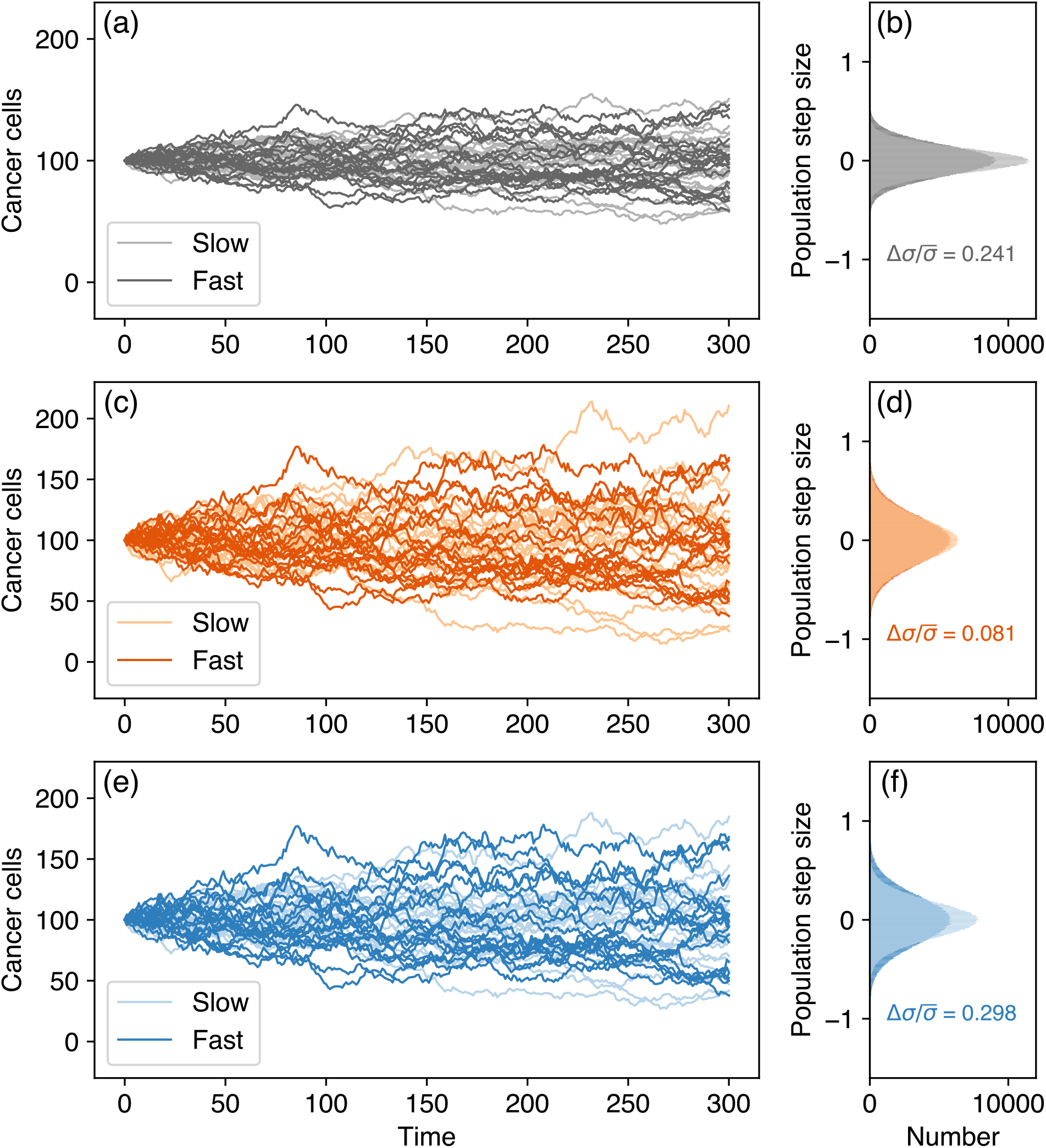
Visualization of contributions from only the diffusion term in Eq. S1.6 (assuming ***a***(***x***) = 0 and V = 25) for (a) and (b) no treatment, (c) and (d) trait-independent treatment with *m* = 1 d^-1^, (e) and (f) trait-dependent treatment with *δ* = 2. The left column shows the time series for 20 replicates. The right column visualizes the population step sizes taken in the simulation (numerical solver evaluation intervals dt = 0.01, plotting time interval 100dt). We use the normalized difference of the standard deviation of the slow and fast subpopulations 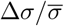 to characterize the different widths of the step size distributions. Large values indicate that the changes of the slow subpopulation are on average smaller than the changes of the fast subpopulation.

**Figure S4.**
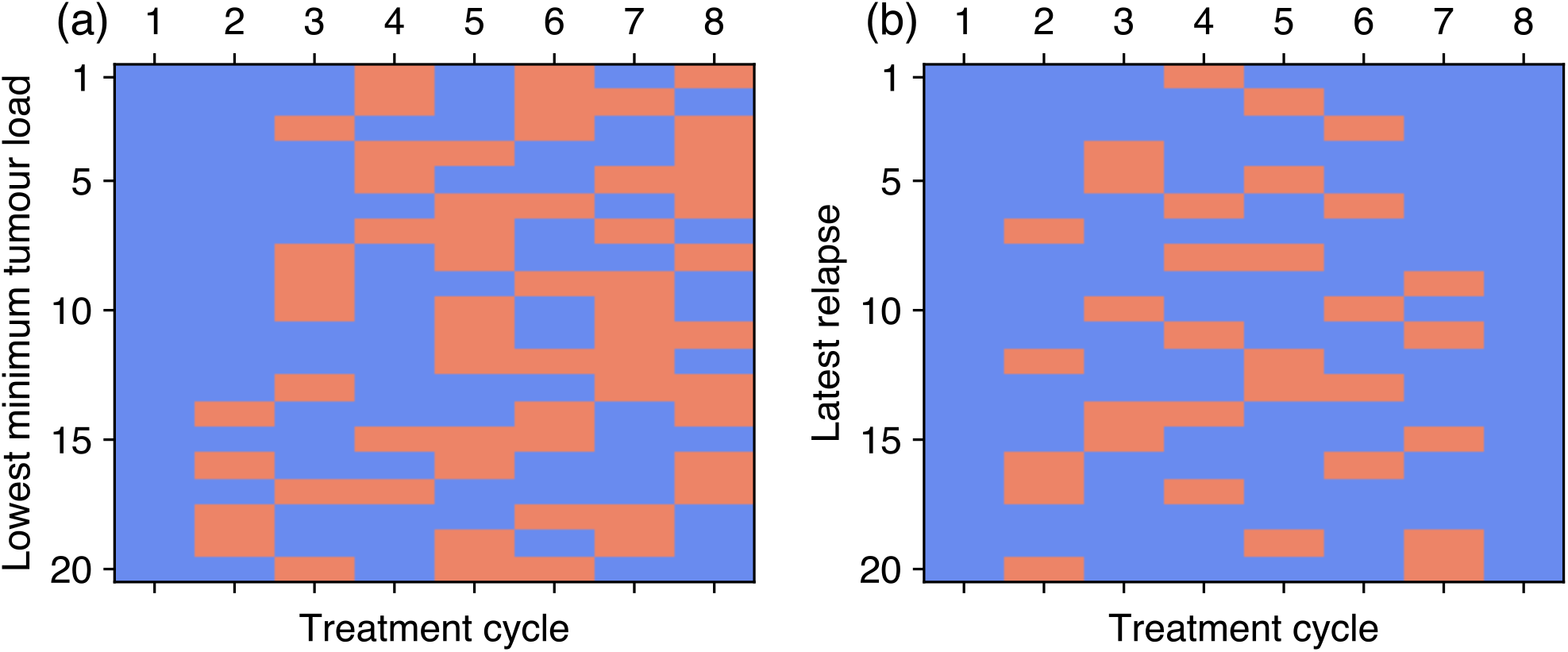
The best-ranked predefined treatment patterns that either (a) result in the lowest minimum tumour load during treatment or (b) reach the tumour load at treatment initiation the latest. Best sequences are at the top, trait-dependent treatment type intervals are blue, trait-independent treatment type intervals are orange. We allowed for 8 different treatment intervals and investigated all 256 combinations.

**Figure S5.**
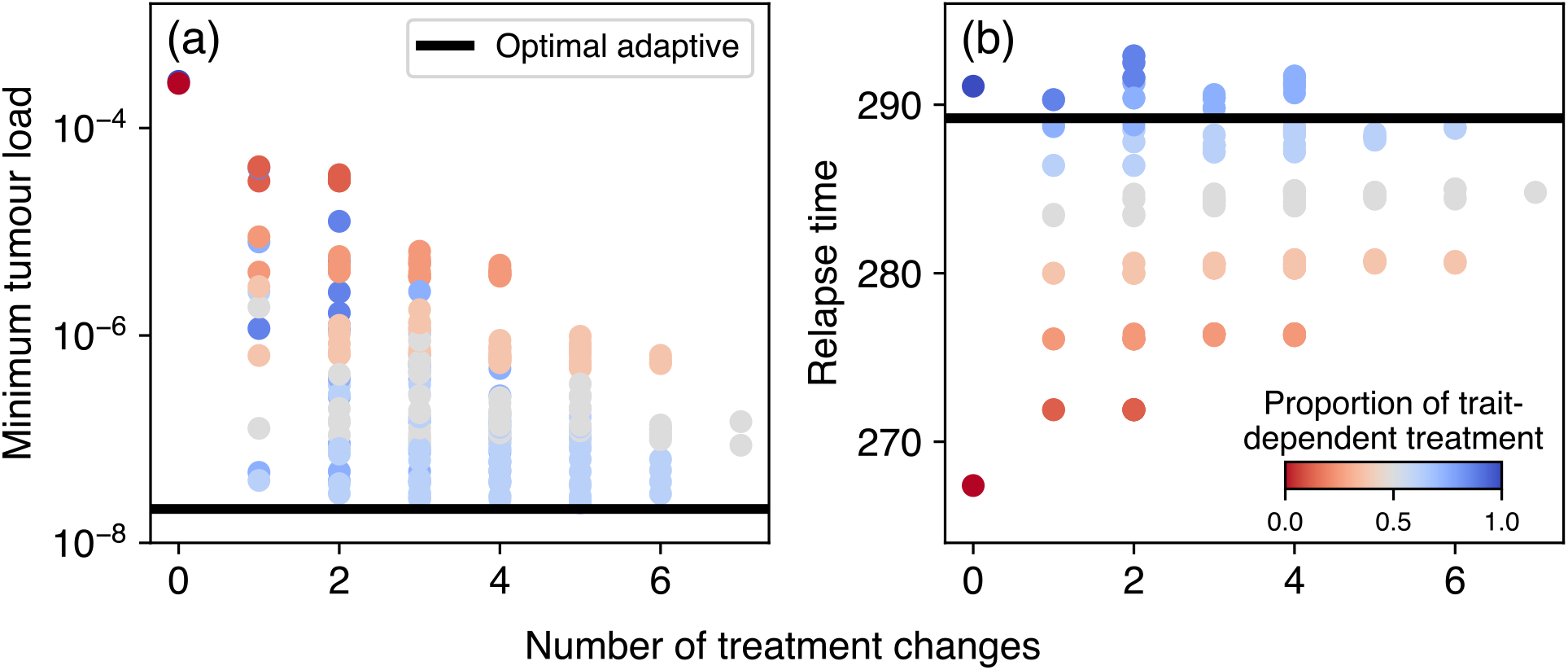
Performance of the predefined treatment scheme for the two treatment goals of (a) minimum tumour load during treatment and (b) relapse time, defined here as the time when the tumour load during the relapse phase exceeds the tumour load at treatment initiation. A maximum of 7 treatment alterations are possible. The blue-to-red colour gradient indicates the proportion of trait-dependent treatment type in every treatment pattern. Note that Fig. 5 shows the correlation of minimum tumour load and relapse time.

**Figure S6.**
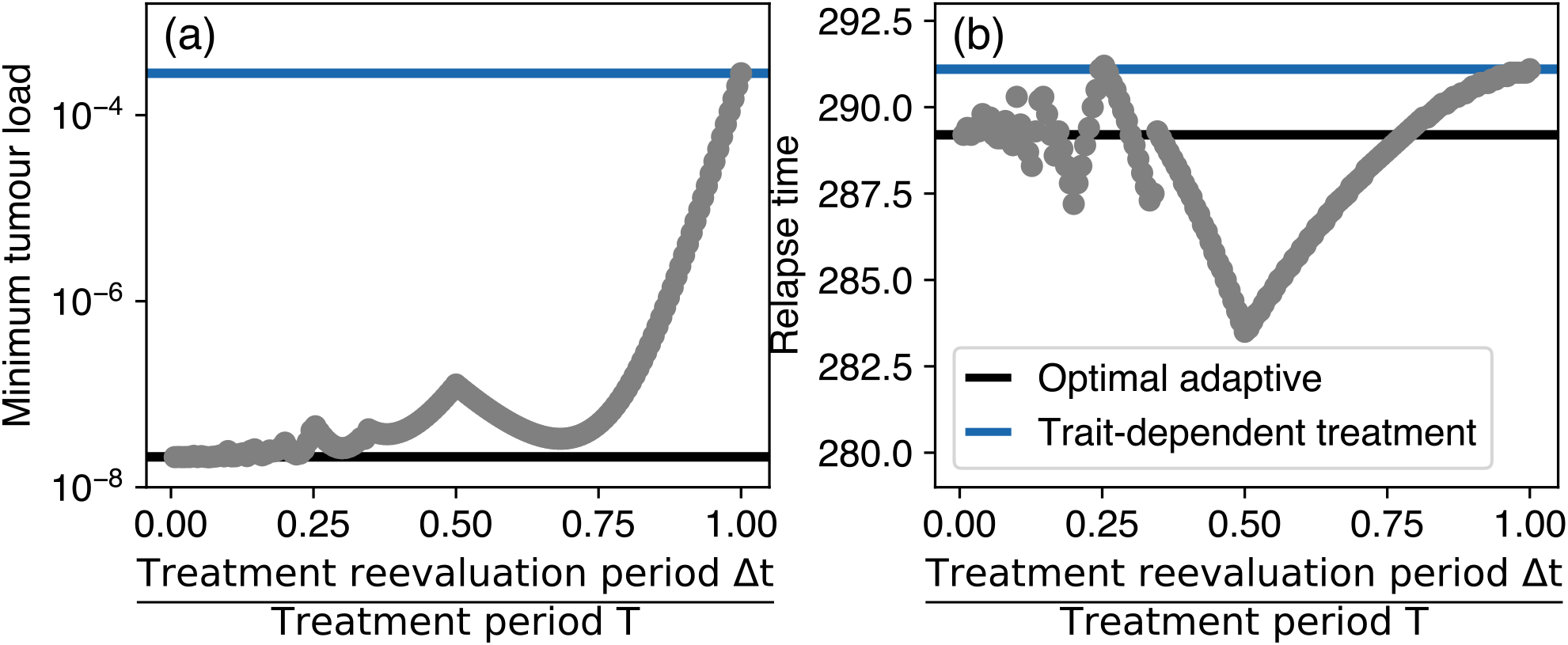
Performance of the realistic adaptive scheme for different reevaluation periods Δ*t* ranges between the optimal adaptive and the purely trait-dependent treatment scheme. Δ*t/T* → 0 corresponds to the optimal adaptive treatment, whereas Δ*t/T* ≥ 1 results in only trait-dependent treatment. The discontinuities arise at reevaluation periods where the number of possible treatment alterations changes. Note that Fig. 5 shows the correlation of minimum tumour load and relapse time.

**Figure S7.**
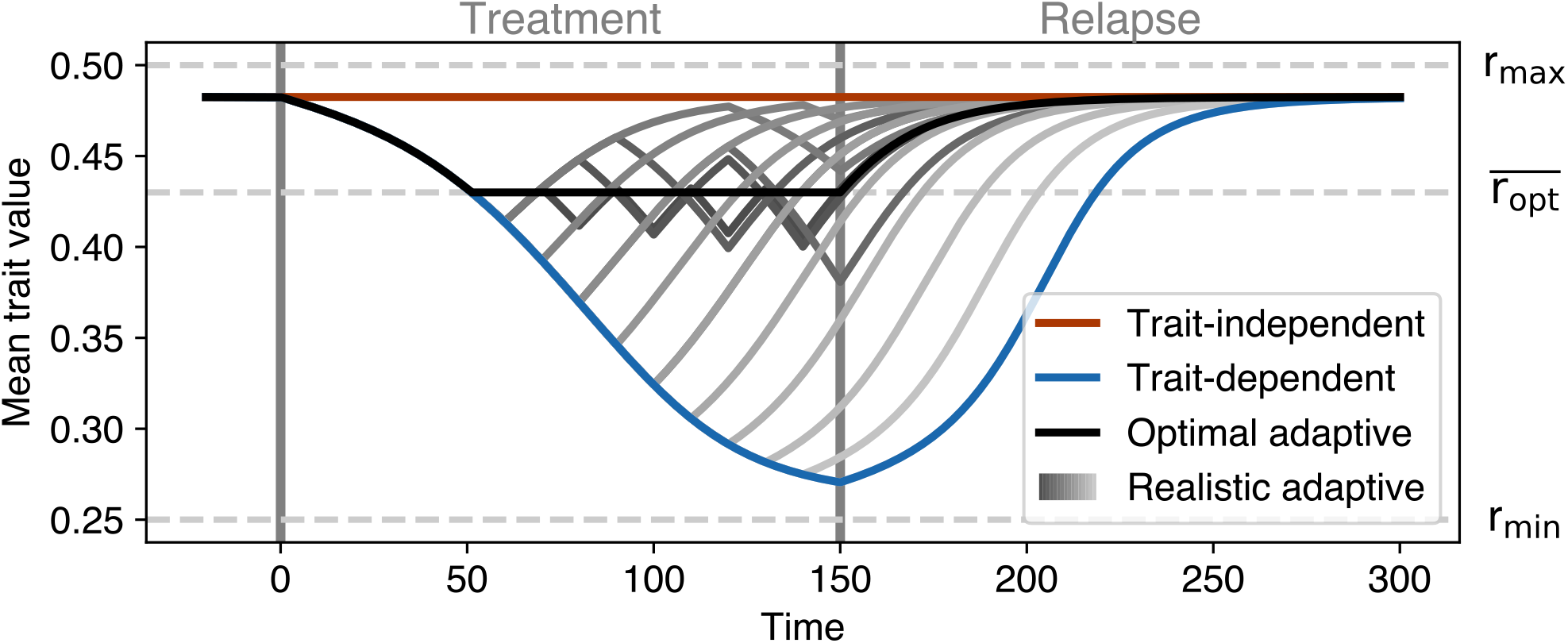
Time series of the mean cancer cell population growth rate for the different treatment schemes. The grey lines correspond to the realistic adaptive scheme with lighter lines showing larger Δ*t*, the difference between them is 10 time units. The optimal adaptive scheme tracks the mean grown rate 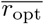 (Eq. 4) where the cancer cell mortality exerted by trait-dependent and trait-independent treatment is equal. The realistic adaptive scheme aims to track 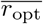 and thus oscillates around it.

